# Use of relevancy and complementary information for discriminatory gene selection from high-dimensional cancer data

**DOI:** 10.1101/2020.02.25.964304

**Authors:** Md Nazmul Haque, Sadia Sharmin, Amin Ahsan Ali, Abu Ashfaqur Sajib, Mohammad Shoyaib

## Abstract

With the advent of high-throughput technologies, life sciences are generating a huge amount of biomolecular data. Global gene expression profiles provide a snapshot of all the genes that are transcribed or not in a cell or in a tissue at a particular moment under a particular condition. The high-dimensionality of such gene expression data (*i.e.*, very large number of features/genes analyzed in relatively much less number of samples) makes it difficult to identify the key genes (biomarkers) that are truly and more significantly attributing to a particular phenotype or condition, such as cancer or disease, *de novo*. With the increase in the number of genes, simple feature selection methods show poor performance for both selecting the effective and informative features and capturing biological information. Addressing these issues, here we propose Mutual information based Gene Selection method (*MGS*) for selecting informative genes and two ranking methods based on frequency (*MGS*_*f*_) and Random Forest (*MGS*_*rf*_) for ranking the selected genes. We tested our methods on four real gene expression datasets derived from different studies on cancerous and normal samples. Our methods obtained better classification rate with the datasets compared to recently reported methods. Our methods could also detect the key relevant pathways with a causal relationship to the phenotype.

## Introduction

Genes are the physical and functional units of hereditary genetic information. The activity and/or expression level of a gene affects the synthesis of downstream proteins that dictate the functionality of a cell. Therefore, the properties as well as the expression levels of a particular set of genes are responsible for a particular phenotype such as disease or tissue morphology. Those genes which are able to differentiate between different states (such as normal vs diseased, quiescent vs proliferating, adult vs stem cells, etc.) of cells are called informative genes or biomarkers (a measurable indicator of a particular state). Identification of these informative genes is very important for elucidating developmental and disease mechanisms, disease diagnosis, drug development, etc. Especially, for different cancer diseases, these informative genes may be invaluable for the improvement of diagnosis, prognosis, and treatment.

Usually, studies to generate cancer specific gene expression profiles comprise a small number of control and patient samples in comparison to tens of thousands of genes (high dimensionality of the data) in each sample where only a few numbers of genes are responsible for a disease. From a large set of genes, identification of a subset that is differently expressed in cancerous cells compared to the normal ones, is a challenging task and is considered as NP hard or NP-complete [1]. Therefore, the feature/gene selection methods can be a useful way to identify a subset of genes relevant to particular cancer for better diagnosis and treatment. In this paper, we use the terms “gene” and “feature” interchangeably.

In bioinformatics, several gene selection methods have been proposed, particularly for cancer data classification [2–4]. “Wrapper”and “Filter”are two popular categories of feature selection methods [5] where wrapper methods are classifier dependent and filter methods are classifier independent and their performance mainly depends on the selection of a criterion. Wrapper based methods select the most discriminant subset of features by minimizing the prediction error of a particular classifier [6]. Support Vector Machine based on the Recursive Feature Elimination (SVM-RFE) [2] is considered to be one of the best performing wrapper methods. It ranks the genes using SVM and selects the important genes combining with the recursive feature elimination strategy. Different variants of SVM-RFE have also been proposed [7, 8]. Although wrapper based feature selection methods provide a better performance, these methods become computationally expensive when the feature size grows. Moreover, these methods are classifier dependent and may not provide the optimal solution for other classifiers [9]. For example, the result of the wrapper method using SVM varies from the result of using random forest (RF). To solve the aforementioned problem, a hybrid filter-wrapper method Information Guided Interactive Search (IGIS) [10] was proposed to select the best genes based on Mutual Information, and the genes were ranked using joint mutual information. However, this method selected more genes than the wrapper or hybrid algorithms. To solve the limitations of IGIS, improved Interaction information-Guided Incremental Selection (IGIS+) [11] was proposed where the first gene is selected based on the highest accuracy and utilizes Cohen’s *d* test to add a new gene into the selected gene set. One major limitation is that it uses several handcrafted thresholds for Cohen’s *d* effect.

Compared to the wrapper methods, filter based methods are more popular as these can assess the property of features without being dependent on any particular classifier. Filter methods select a subset of features based on some criteria that can be evaluated on the dataset itself. Different criteria that filter methods use are: correlation coefficient [12], t-statistics [13], and mutual information [14, 15]. Among these, MI based methods are the most popular for feature selection due to their strong theoretical background and ability to capture non-linear dependencies between features. MI based methods have also been applied for gene expression data analysis [16]. One of the earliest works used Minimum Redundancy Maximum Relevance (MRMR) [3]. In this method, the authors select each gene incrementally which holds the highest discriminatory power (relevancy) with the target class (control/cancer) and lowest dependency (redundancy) with other selected genes. Relevance with the class variable is calculated using MI between a gene and class variable or F-statistics while redundancy among the genes is calculated using pair-wise MI or Pearson correlation coefficient for discrete and continuous data, respectively. MRMR based methods have also been adopted for gene selection using temporal gene expression data [17].

Feature selection methods that use the aforementioned criteria are often posed as an optimization problem where the goal is to select the feature subset that optimizes a cost function. Generally, this cost function is constructed using one of the above criteria. Apart from the aforementioned wrapper and hybrid methods, there exist few popular evolutionary or bio-inspired algorithms to search for the optimal set of features. Almugren et al. in [18] provide an extensive review of the bio-inspited wrapper and hybrid methods. Alshamlan et. al. proposed a hybrid artificial bee colony as well as a genetic bee colony optimization method that uses mRMR criterion [19, 20]. El Akadi et al. [21] proposed a genetic algorithm based on mRMR criteria. In these methods, mRMR criterion is used to filter noise and redundant genes in the high-dimensional microarray data and then the bio-inspired algorithm uses the classifier accuracy as a fitness function to select the highly discriminating genes. Most bio-inspired algorithms are local searches with random restart or population based methods. However, these algorithms still can get stuck at a local optimum. In order to solve the optimization problems globally, several selection methods were attempted [22]. These methods incorporate parallel search strategies based on semi-definite programming (SDP) or quadratic programming that can find the feature subset in polynomial time [23].

Recently, deep learning based methods have provided better accuracy in different classification problems such as image or audio classification [24]. These deep learning based architectures have also been proposed for classification problems using gene expression data [4, 25]. One of the most recent works based on deep learning was proposed by Ding and Peng [4]. The authors developed a new model namely Forest Deep Neural Network (fDNN) that incorporates deep neural network (DNN) with random forest (RF) to solve the problem of learning from small sample data having a large number of genes. RF is used to reduce the dimension of these datasets by detecting the important genes in a supervised manner [26]. This new feature representation was then fed into DNN to predict the outcomes. However, this method does not make use of the main advantage of deep learning in solving classification problems, which is automatic feature extraction. On the other hand, using a neural network as a black box to extract new features from gene expression data also reduces the interpretability of the classifier which is important in studies such as cancer classification.

Since MI based filter methods do not extract new features and thus are more interpretable, parallel to the development in Deep learning, there has been a lot of effort to better approximate MI measures such as relevancy and redundancy. New Information theoretic measures such as complementary information, the additional information that a gene has about the class, which is not found in the already selected subset of genes have been proposed [15, 27]. These methods attempt to estimate the joint mutual information of a feature subset with the class. In mDSM [15], the authors showed that during the calculation of MI for finite samples, there exist some errors (bias) for all the three terms namely relevancy, redundancy and complementary. Moreover, for selecting a feature, they proposed to use *χ*^2^ statistics by showing that all three terms follow *χ*^2^ distribution. Moreover, even though it has few good characteristics, by incorporating the term *redundancy* in gene expression data, informative genes might be discarded [11]. Another issue with gene selection for cancer classification, in contrast to traditional feature selection methods in machine learning, is that the set of genes selected should be biologically relevant to the disease under study. Although filter methods can produce a subset of genes that may be highly accurate in classifying cancer, the literature on filter methods rarely discusses the biological relevance of the selected genes [5].

To solve the aforementioned problems, we propose a new MI based filter method namely Mutual information based Gene Selection (*MGS*) that achieves better classification performance with high dimensional biological data. The main contributions of this paper are as follows: first, a gene selection technique is proposed for identifying discriminating genes based on their relevancy and complementary information. Second, a statistical test is used to select genes without a handcrafted threshold. Third, two ranking techniques are proposed for the selection of informative genes that are used in cancer classification. Finally, the proposed methods select the relevant genes associated with a particular type of cancer.

## Materials and methods

In this section, we firstly describe the datasets that are used in our experiment and then discuss the proposed gene selection method. It selects some candidate genes and then ranks the genes based on one of the two proposed ranking criteria. Finally, using the selected top *η* genes, classification and biological interpretations are then performed.

### Dataset description

We used four different gene expression datasets GDS3341 [28], GDS3610 [29], GDS4824 [30] and GSE106291 [31] retrieved from the Gene Expression Omnibus (GEO) database [32] at the National Center for Biotechnology Information (https://www.ncbi.nlm.nih.gov). GDS3341 and GDS3610 are independent experimental datasets generated from nasopharyngeal carcinoma (NPC) tissue samples. GDS4824 contains the gene expression data of malignant and benign prostate cancer tissues. GSE106291 contains the RNA-seq expression profiles of acute myeloid leukemia patients for the prediction of resistance to induction treatment. The description of datasets are given in Table 1. We used two different global gene expression datasets of nasopharyngeal carcinoma tissues (GDS3341 and GDS3610) as built-in controls in the study to assess the coherence and performance of our proposed methods. Expression data of multiple probes for the same gene were merged. All these datasets contained much less number of samples compared to the number of genes (Table 1).

**Table 1.** Summary of the datasets used in this study.

| Dataset ID | Total samples | Control samples | Cancer samples | Features(genes) |
| --- | --- | --- | --- | --- |
| GDS3341 | 41 | 10 | 31 | 30865 |
| GDS3610 | 28 | 3 | 25 | 14126 |
| GDS4824 | 21 | 8 | 13 | 30872 |
| GSE106291 | 235 | 71 | 164 | 21403 |

### Overview of gene selection and validation process

In this paper, we propose an MI based Gene Selection (*MGS*) method for the selection of an informative gene subset that provides both better classification accuracy and contains biologically relevant information for cancer gene identification. The overall process of *MGS* is shown in Fig 1 where we first identify the informative gene subset (Fig 1A) and then use the top *η* genes from that subset for classification (Fig 1B). The following subsections describe our method with further details.

**Fig 1.**
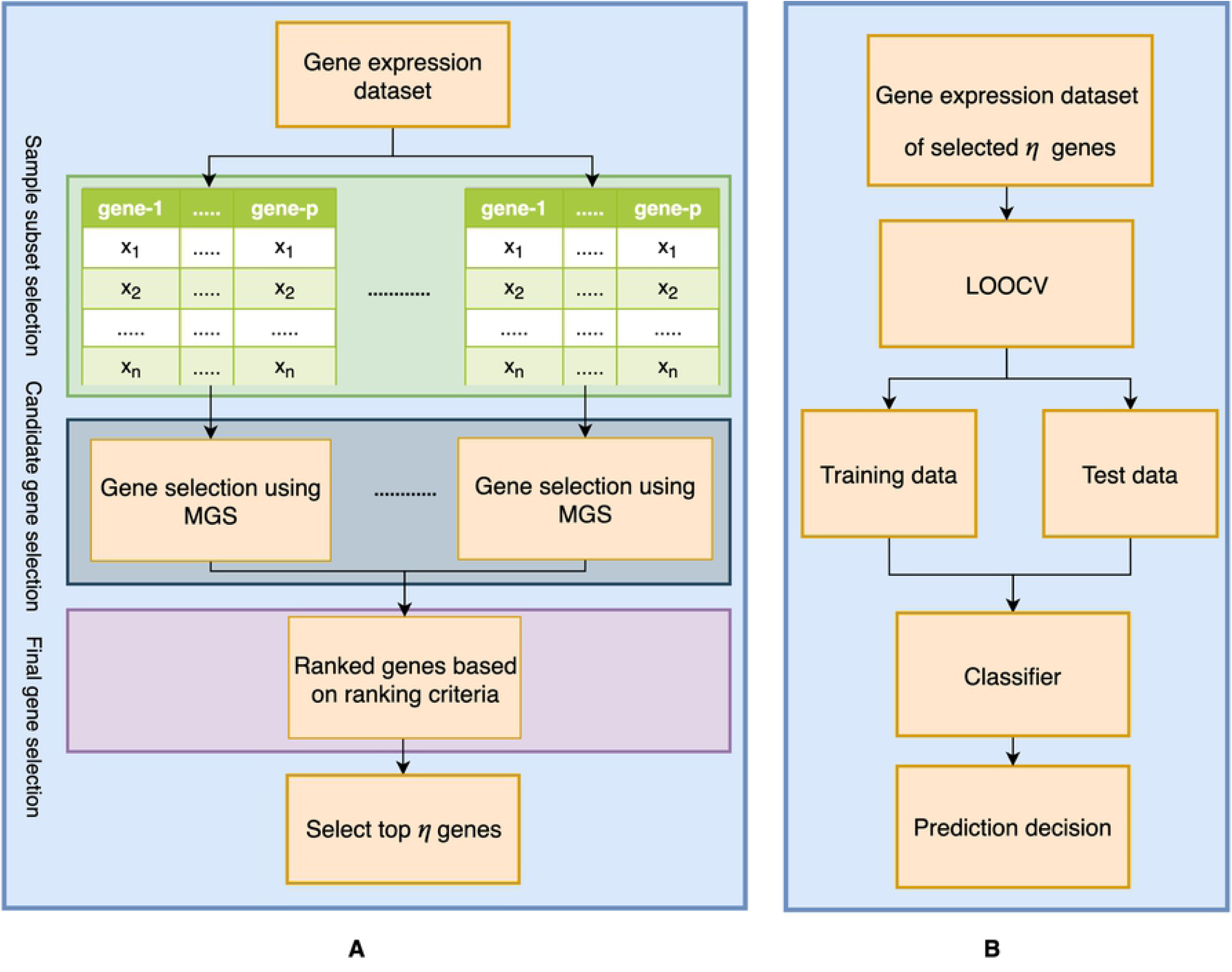
Overall process of the proposed method. (A) Gene selection. (B) Classification

### Gene subset selection

For the identification of a gene subset, in this paper, we propose to use a filter based gene selection method that approximates the joint MI with respect to the class variable. In order to identify an informative gene subset, we first subdivide the given gene expression dataset into *K* sets. This can be done through a cross validation process when we have a large number of samples (*n*). However, when *n* is small, Leave-one-out cross validation (LOOCV) is proposed here to apply. Since, in this study, the main focus was the identification and classification of genes in datasets with a small number of samples having a very large number of genes, we applied LOOCV. In *MGS*, we incorporate a variant of *mDSM* [15] by modifying the selection criteria so that it can identify biologically relevant genes for a class. The accumulation of all genes identified by *MGS* from *K* different subsets is defined here as candidate genes (*G*_*SC*_). The final selected gene subset (*G*_*S*_) is then obtained by ranking the candidate genes (*G*_*SC*_). Two ranking criteria namely *MGS* frequency-based ranking (*MGS*_*f*_) and *MGS* Random Forest (RF) based ranking *MGS*_*rf*_) are proposed here to select the top *η* genes as biomarkers for cancer classification.

#### Candidate gene selection

To measure how much information a particular gene expression provides for the identification of cancer data, we calculate MI between an expression value of a gene *g*_*i*_ and the class variable *C*. This MI represents the relevancy of a gene that reveals the degree of importance of that gene in cancer data classification. Note that, before calculating the MI, the gene expression data is quantized which is necessary for noise reduction and data simplification and thus result in maximizing the relevancy of a gene to the target class *C*. For calculating the relevance between *g*_*i*_ and *C*, MI is calculated using Eq. 1.

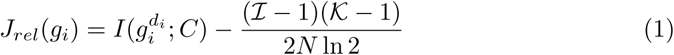

where, 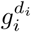 denotes gene *g*_*i*_ with *d*_*i*_ discretization levels. For each gene *g*_*i*_, the minimum discretization levels *d*_*i*_ is chosen for which *J*_*rel*_(*g*_*i*_) is greater than its *χ*^2^ critical value 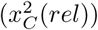. This test helps to determine whether the gene is significantly relevant or not and can be done because it can be shown that the relevancy follows *χ*^2^ distribution with (ℐ − 1)(𝒦 − 1) degrees of freedom. Here, ℐ, 𝒦 and *N* represent the quantization levels of gene *g*_*i*_, the total number of classes in *C* and the total number of samples respectively. The genes which do not satisfy the *χ*^2^ critical value are discarded considering these genes are not related to *C*. All the genes selected through this process are now ranked in descending order based on the relevancy. From this ranking, a selection method is followed to get a subset of informative genes. As the top ranked gene is considered to be the most important, we include it to the candidate gene subset *G*_*SC*_ at first. Now, the second ranked one is evaluated for selection based on its score calculated using Eq. 2.

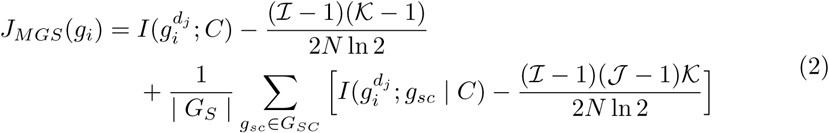

here, along with relevancy, the complementary information 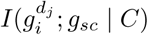 of a new gene is also incorporated. The complementary information 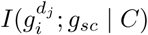 due to *g*_*i*_ for an already selected gene *g*_*sc*_ reveals the dependency among those genes while identifying the class variable *C*. Here, 𝒥 represents the quantization levels of each gene *g*_*sc*_ in *G*_*SC*_. In Eq. 2, the bias correction is also incorporated for calculating relevancy and complementary information. While calculating the value of *J*_*MGS*_, the quantization level (*d*_*j*_) of the *g*_*i*_ is also shifted by a small amount (±*δ*). This is because a small shifting of quantization may increase the value of *J*_*MGS*_ and this new quantization value is chosen dynamically considering the dependency among the genes. Now, for a particular gene (*g*_*i*_), if the value of *J*_*MGS*_ is larger than the *χ*^2^ critical value 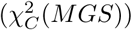, then it is placed into the selected gene subset. It means when the relevancy and complementary information of a *g*_*i*_ is significant, then it is selected otherwise discarded. So, finding genes that maximize *J*_*MGS*_ indicates the genes which are strongly relevant with the class *C* with greater complementary information will be adopted to the selected subset throughout this process.

It is noteworthy to mention here that there may exist a group of genes that share similar information and thus their expression values which may even make them redundant. However, if they have complementary information about the class, it is necessary to incorporate that gene into the selected subset even though it is redundant. Such incorporation of the redundant genes is logical because usually a set of genes contribute mutually for a particular task in our body and these genes may share a similar expression profile. Hence it is required not to consider the redundancy in criteria of gene selection. The biological importance of such exclusion is also shown experimentally in the result and discussion section.

#### Final gene selection

The same subset of genes is not always selected during the selection of genes by *MGS* at each iteration of LOOCV. In the final gene selection step, we aggregate all the candidate gene subsets (*G*_*SC*_) from the candidate gene selection step and find the union of these subsets, *G*_*S*_. Afterward, these genes in *G*_*S*_ are ranked using one of the following two ranking criteria.

- *MGS*_*f*_ : This ranking is performed based on the following assumption. *Assumption*: The genes which are selected in every iterations are likely to have more discriminating power and biological significance. To quantify the *Assumption*, we compute the relative frequency of every selected gene, *S*_*i*_ in *G*_*S*_ using Eq. 3.

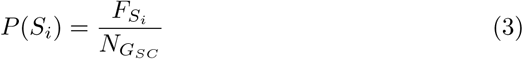

here, 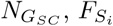 and *P*(*S*_*i*_) are the total number of candidate subsets, frequency of the selected gene *S*_*i*_ and the relative frequency of gene *S*_*i*_ respectively. For example, we have two candidate gene subsets from candidate gene selection step, *L*_1_ = {*g*_1_, *g*_3_, *g*_4_, *g*_5_, *g*_6_} and *L*_2_ = {*g*_1_, *g*_2_, *g*_4_, *g*_6_}. Here, the unique genes are *G*_*S*_ = {*g*_1_, *g*_2_, *g*_3_, *g*_4_, *g*_5_, *g*_6_} and the frequencies of these unique genes are *F* = 2, 1, 1, 2, 1, 2 respectively. So, the relative frequency is *P*(*S*_*i*_) = 2/2, 1/2, 1/2, 2/2, 1/2, 2/2. Thus, based on the *P*(*S*_*i*_), ranked genes are *G*_*S*_ = *g*_1_, *g*_4_, *g*_6_, *g*_2_, *g*_3_, *g*_5_.
- *MGS*_*rf*_ : Informative genes have the ability to split the control and cancer samples into two groups. To find the more informative genes, we need to rank the candidate genes. In order to rank these genes, it is necessary to measure how much information a gene contains. To measure the information content of a gene, we can use Information Gain (IG) criterion. IG is used in decision trees to select features that reduces the entropy of the data most by splitting data into two groups (called the the left and right child in a decision tree). We use weighted IG derived in Eq. 4.

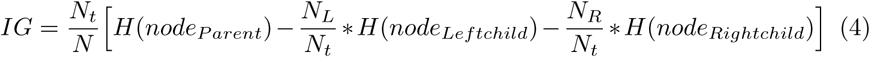

where, *N*_*t*_ is the number of samples at the current (parent) node, *N* is the total number of samples, *N*_*L*_ is the number of samples in the left child, and *N*_*R*_ is the number of samples in the right child. *H*(*node*) is the entropy at the node. The entropy is calculated using Eq. 5.

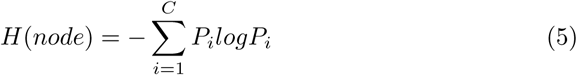

This weighted IG is commonly used in Decision Tree (DT) [26], where each node in a DT contains a gene with its corresponding weighted IG. Besides, to make the weighted IG more robust, we use *M* number of DTs to construct a Random Forest and take the average of IGs for each gene *g*_*j*_ ∈ *G*_*S*_ using Eq. 6.

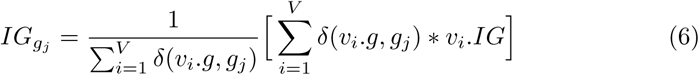

here, *V* = {*v*_*i*_, *v*_*i*+1_,..,*v*_*k*_} = {(*g*_*i*_, *IG*_*i*_), (*g*_*i*+1_, *IG*_*i*+1_),..,(*g*_*k*_, *IG*_*k*_)} and *k* is the total number of nodes in the random forest. That is, for each node of the random forest, we store the corresponding gene and its weighted IG in *V*. *δ*(*v*_*i*_.*g, g*_*j*_) = 1 if *v*_*i*_.*g* = *g*_*j*_, and 0, otherwise (Kronecker function).
This average score can be used as the importance score of each gene. In our case, this importance score represents how important a particular gene is to explain the target class. Finally, based on the importance score, the genes from *G*_*S*_ are ranked in descending order.

### Classification

In this stage, as shown in Fig 1B, only selected genes from the previous step are used in the train and test data to fit the classifiers and predict the outcome. Due to a limited number of samples in each data set, we employ LOOCV to partition all the data samples into training and testing sets. For example, a dataset having *n* number of samples, we used (*n* − 1) samples for training and the *n*^*th*^ sample for testing. After passing the training data to *MGS*, we get candidate informative genes. This is repeated *n* times and passing the selected candidate gene to *MGS*_*f*_ and *MGS*_*rf*_ for finding the ranked genes. And finally, from the ranked genes, we take top *η* genes as biomarkers and calculate the performance metrics.

To assess the performance of a gene selection method, we consider two performance metrics *accuracy* and Area Under the Receiver Operating Characteristic Curve (*AUROC*). *Accuracy* is the percentage of samples that are predicted to the true class. *AUROC* represents degree or measure of separability between classes and it can be used when the dataset is highly imbalanced and the number of samples is less than the number of genes. *ROC* is a probability curve of a classifier at various thresholds. It plots curve based on the true positive rate (TPR) and false positive rate (FPR) represented in Eq. 7 and 8.

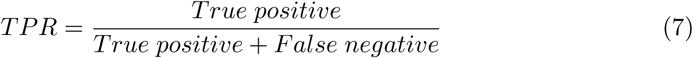

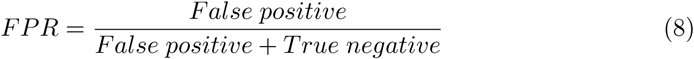

here, “True positive” and “True negative” are the numbers of positive and negative samples that are correctly classified. “False positive” are the numbers of negative-class samples misclassified as the positive class, and “False negative” are the numbers of positive-class samples misclassified as the negative class. To compute the points in a *ROC* curve, *AUROC* computes an aggregate measure of various thresholds. For our experiments, the reported results are calculated by taking the average over the LOOCV process for these two metrics.

### Biological interpretation of the selected genes

We used NetworkAnalyst [33] to interpret the biological significance of the selected gene. NetworkAnalyst is a bioinformatics platform to interpret gene expression data within the context of protein-protein interaction (PPI) networks. We used top *η* selected genes for each dataset determined by our proposed and the previously described methods as input in NetworkAnalyst. Since the type of the cancer samples in the datasets was known, we assessed the performance of the compared methods based on their abilities to identify the key pathways affected in the corresponding cancer types.

## Results and Discussion

We compared the performances of our proposed ranking methods (*MGS*_*f*_ and *MGS*_*rf*_) to other renowned methods-*RF, fDNN, IGIS*+, and *mDSM*. As *mDSM* is a gene selection method, so we incorporate our frequency and RF based gene ranking methods (*mDSM*_*f*_ and *mDSM*_*rf*_) for comparison purpose. Note that, for a fair comparison, we followed the same training and testing protocol for all the datasets. For *RF, fDNN* and *MGS*_*rf*_ (where random forest is used) we used 300 decision trees. We evaluated the performance of these methods using SVM (linear kernel) and RF classifiers. These classifiers are implemented in Python with packages Scikit-learn [34].

In this experiment, we applied the aforementioned methods on four gene expression datasets. In this section, we first discuss the performance of all methods in terms of *accuracy* and *AUROC* and then, provide the biological interpretation selecting top *η* (= 10) genes. In situations where feature selection method (*IGIS*+, *mDSM*_*f*_, *mDSM*_*rf*_) had selected less than 10 genes, we used only these genes for our analysis. We also discuss the performance of a different number of top *η* genes for measuring the robustness of our method.

### Classification performance

Table 2 summarizes the comparative results of the proposed methods along with the existing methods on four datasets as mentioned before. Analyzing the table, it becomes evident that our proposed methods (*MGS*_*f*_, *MGS*_*rf*_) performed better than than the other methods (*RF, fDNN, IGIS*+, *mDSM*_*f*_ and *mDSM*_*rf*_) in classification results in terms of both *accuracy* and *AUROC* (Table 2), which indicate that our methods selected more informative genes. In the case of dataset GDS3341 and GDS4824, all methods except *RF* were able to perfectly differentiate the control and cancer disease for both *SV M* and *RF* classifiers. The small number of samples compared to a large number of genes may be the reason behind the relatively poor performance of *RF*. However, even though other methods performed well for selecting distinguishable genes, all the genes were not biologically informative (discuss in the next subsection). For the dataset of GDS3610 and GSE106291, *MGS*_*f*_ and *MGS*_*rf*_ methods achieved much better *accuracy* and *AUROC* compared to the other methods. The performance of *MGS*_*f*_ was better to *RF, fDNN, IGIS*+, *mDSM*_*f*_ and *mDSM*_*rf*_ in most instances inspite of having imbalanced dataset and *n* ≪ *p* property. Moreover, *MGS*_*rf*_ unequivocally performed better compared to *MGS*_*f*_.

**Table 2.**
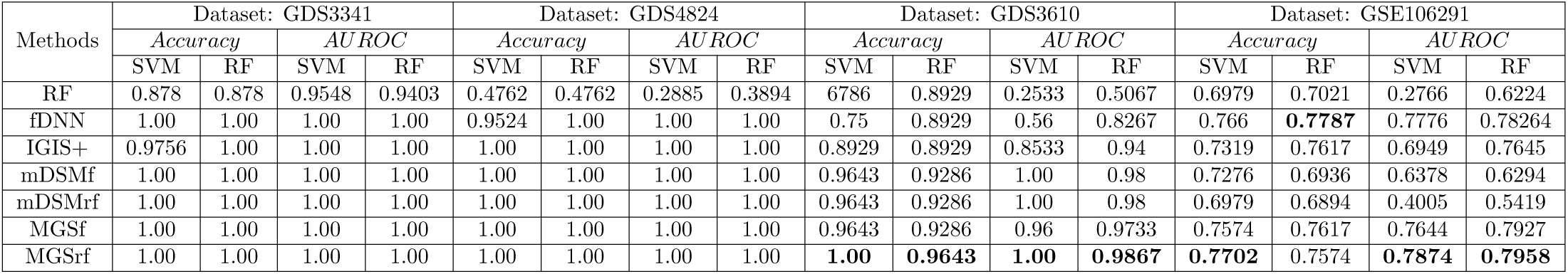
Classification accuracy and AUROC of different methods for GDS3341, GDS4824, GDS3610 and GSE106291 datasets.

It is not always true that the selected genes that have better classification ability are also relevant for a biological process. To examine this, apart from *accuracy* and *AUROC*, we investigated the ability of the top (≤ 10) selected genes in identifying the most relevant pathways in the cancer types used in different datasets. This is described in the next section.

### Biological interpretation

From Table 2 and Table 3, it is evident that *MGS*_*f*_ and *MGS*_*rf*_ not only achieved better *accuracy* and *AUROC* but also performed better in capturing the genes more relevant to the cancer type. For example, Epstein-Barr virus (EBV) is well known to cause nasopharyngeal carcinoma (NPC), which is epithelial cancer prevalent in Southeast Asia [35–37]. GDS3341 and GDS3610 datasets contain NPC samples [28, 29]. Though GDS3341 and GDS3610 are independent datasets, both *MGS*_*rf*_ and *MGS*_*f*_ could detect the genes involved in viral carcinogenesis and Epstein-Barr virus infection (Table 3). We used two different datasets (GDS3341 and GDS3610) on the same cancer type as built-in controls in the study to increase confidence in the experimental results. With both the datasets *MGS*_*rf*_ and *MGS*_*f*_ performed almost equally well. Moreover, the genes selected by *MGS*_*rf*_ performed better than those selected by the *MGS*_*f*_. The other methods (*RF, fDNN, IGIS*+, *mDSM*_*f*_ and *mDSM*_*rf*_) could detect these pathways only with the GDS3610 dataset. In fact, *RF* and *IGIS*+ could detect one of these pathways. The GDS4824 dataset contains gene expression data from prostate cancer samples. Both the *MGS*_*rf*_ and *MGS*_*f*_ detected genes that are involved in prostate cancer. Although the prostate cancer pathway was ranked 6^*th*^ in the detected pathways (based on the FDR values) with the genes selected by the *MGS*_*rf*_ and *MGS*_*f*_, the top ranked pathways (FoxO signaling pathway, colorectal cancer, pancreatic cancer and endometrial cancer) are relevant to cancer as well [38–41]. In fact, unlike nasopharyngeal carcinoma, prostate cancer development involves different pathways. Fork head box O transcription factors (FoxO) regulates multiple cellular processes, including cell cycle arrest, cell death, DNA damage repair, stress resistance, and metabolism [42]. Inactivation of FoxO protein is linked to multiple tumorigenesis including prostate cancer [42–44]. Among the other methods, *fDNN, mDSM*_*f*_ and *mDSM*_*rf*_ could detect the genes associated with prostate cancer, although the rank of the pathway and associated FDR values were less significant.

**Table 3.**
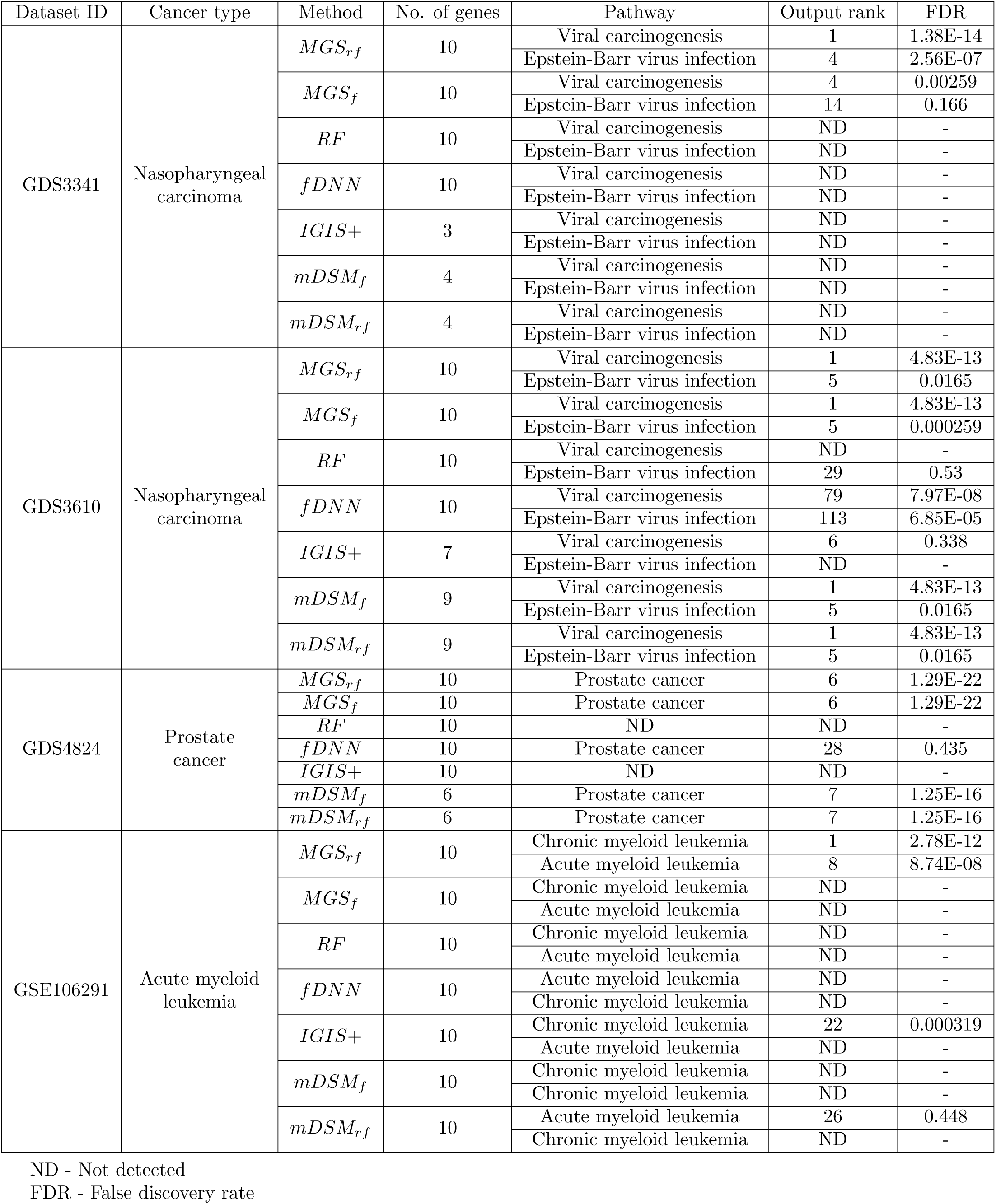
Comparative performance of different methods in identification of relevant biological pathways.

It is well known that, although multiple proteins interact in a network in a cell to attain a particular function, each of these does not play an equally important role. Some proteins in a network are more connected and play a pivotal role in the overall biological process. *MGS*_*rf*_ and *MGS*_*f*_ selected top genes play important roles in pathways relevant to cancer (Fig 2) whereas other methods could not detect any pivotal genes relevant to cancer (Table 3).

**Fig 2.**
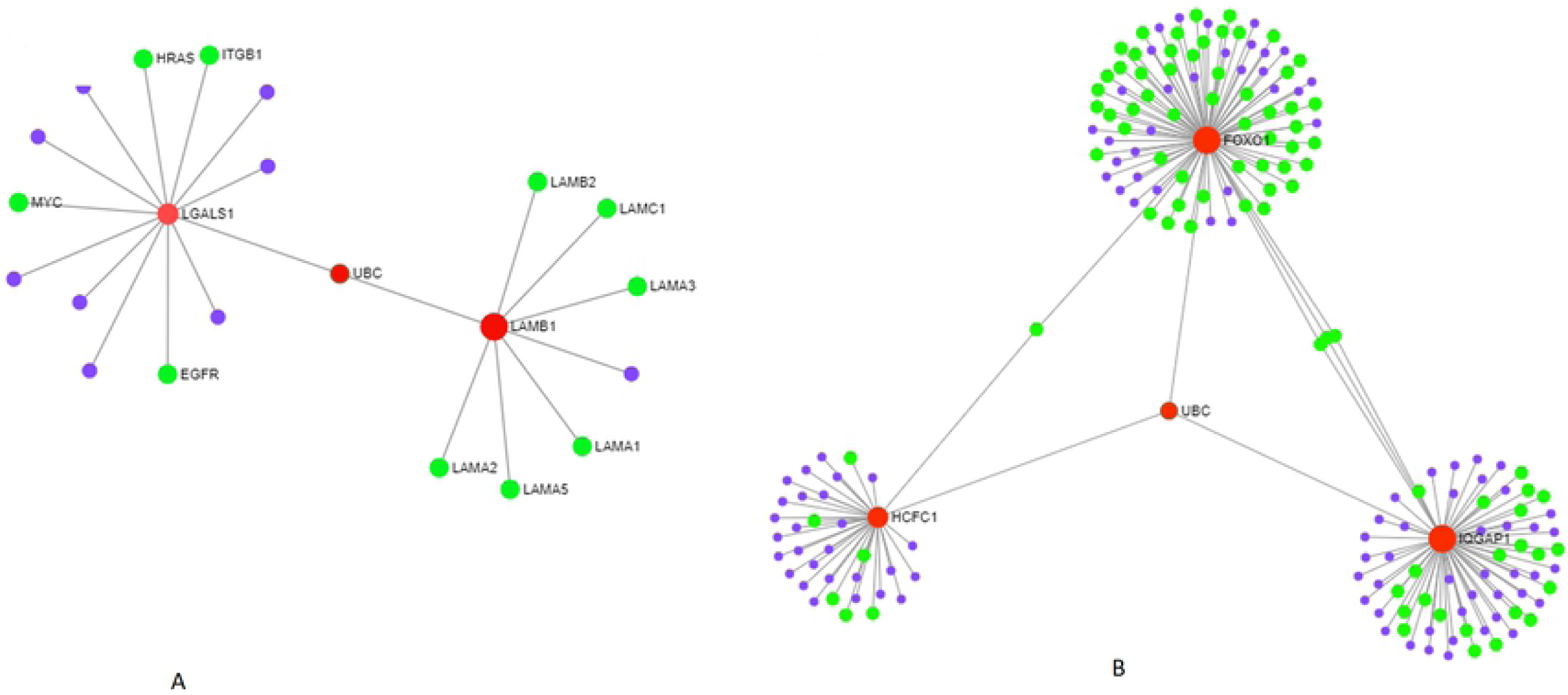
Roles of *MGS*_*rf*_ selected top genes in pathways related to cancer. (A) *LGALS*1 and *LAMB*1 were selected among the top 10 genes from GDS3341 dataset by the *MGS*_*rf*_ method. These (highlighted in red) are part of a sub-network that contains many other proteins (highlighted in green) known to play roles in different cancers [33]. (B) *HCFC*1, *FOXO*1 and *IQGAP*1 were selected among the top 10 genes from GDS4824 dataset by the *MGS*_*rf*_ method. These (highlighted in red) are part of a sub-network that contains many other proteins (highlighted in green) known to play roles in different cancers [33].

It is noteworthy to mention here that the proposed methods perform much better compared to *mDSM* even though they follow a similar methodology. The main difference is the exclusion of redundancy term. However, such avoidance of redundant genes may not be appropriate for selecting genes as genes working together in a pathway may be regulated in a more coordinated fashion than a random set of genes and thus, share a more coherent expression profile [45]. Therefore, *MGS*_*f*_ and *MGS*_*rf*_ do not consider redundancy in Eq. 2 to select new genes. For example, *mDSM* discards a gene *g*_*i*_ if it finds another gene, *s*_*i*_ with similar expression level. But as mentioned earlier, both *g*_*i*_ and *s*_*i*_ may be informative despite being considered “redundant” and may add complementary information for a disease if selected instead. To understand this issue, let us consider an example of two genes named *MAN*1*C*1 and *ARCN*1 in dataset GDS3610 where *MAN*1*C*1 is on the selected list and *ARCN*1 is considered to be on the selected list. As the redundancy value (0.685461) is greater than *χ*^2^ critical value (0.558168), *mDSM* discarded *ARCN*1. However, our methods selected *ARCN*1 in the selected list as it provides complementary information (0.598510). Despite the fact that these genes work in different pathways, both inhibit cancer cell proliferation [46, 47].

### Comparison of performances for different number of genes

We also investigated the performances of the aforementioned methods for a different number of selected genes (*η*) using two metrics *accuracy* and *AUROC* as shown in Figs 3, 4, 5 and 6. Except *RF*, all the methods performed well (Figs 3-6). In the case of GDS3341 and GDS4824 datasets, for a different number of genes, all the gene selection methods classified the samples almost perfectly as shown in Figs 3 and 5. For these two datasets, the expression values of genes are more distinguishable between classes which would be the reason for the almost equal performance of every method. That would be the reason why the performance is not varied with the increasing number of selected genes. For the small and highly imbalanced dataset GDS3610, our methods showed its superiority for a different number of *η* genes (Fig 4). Our methods handled not only small datasets but also imbalanced dataset which is shown in Fig 4B, as *AUROC* is a better metric for imbalanced datasets. We also showed our method’s strength in dataset GSE106291, having comparatively large samples (Fig 6). Here, our methods performed better than others in terms of *accuracy* and *AUROC* over the different *η*, indicating its applicability on gene expression datasets with small and relatively medium sample size.

**Fig 3.**
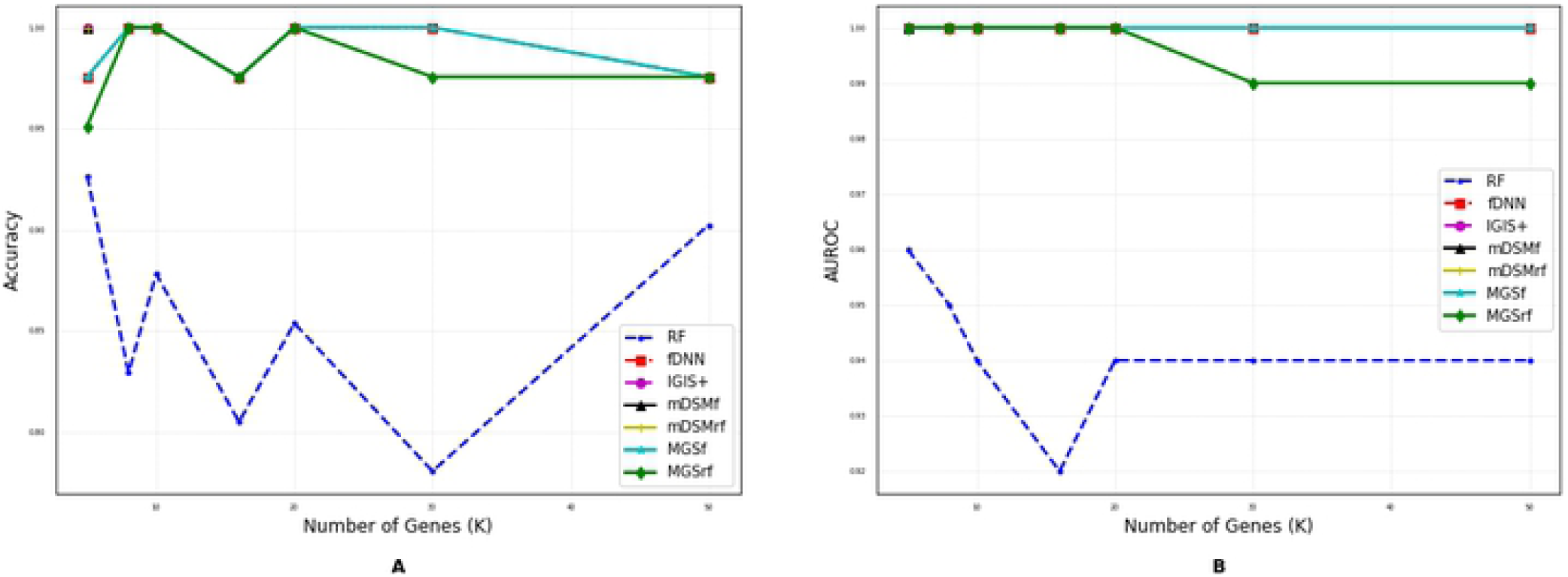
Performance comparison using different number of selected genes for the GDS3341 dataset. (A) Accuracy. (B) AUROC.

**Fig 4.**
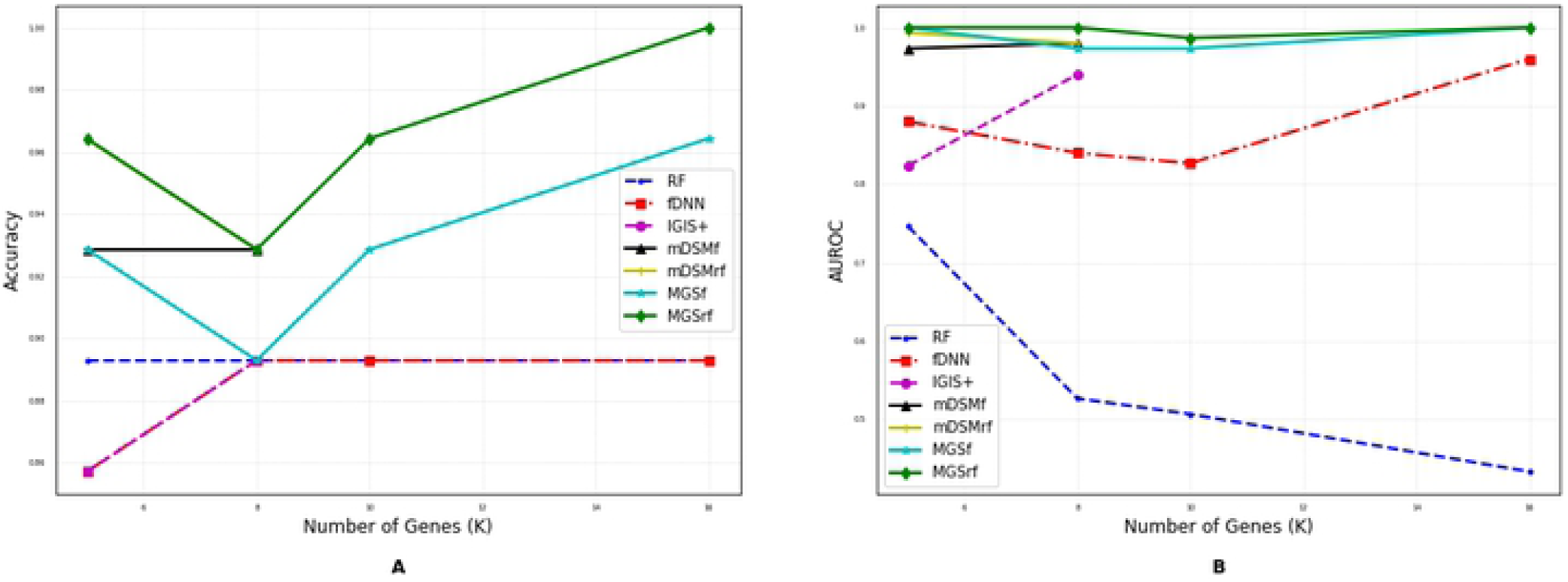
Performance comparison using different number of selected genes for the GDS3610 dataset. (A) Accuracy. (B) AUROC.

**Fig 5.**
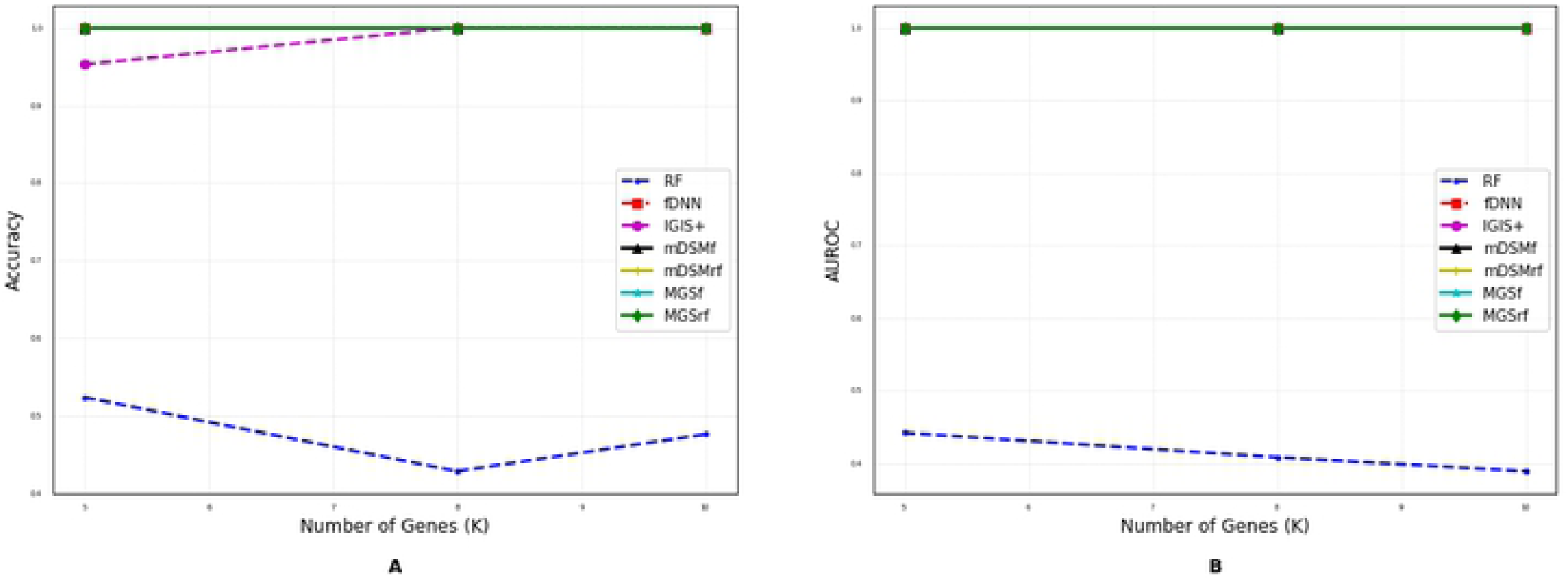
Performance comparison using different number of selected genes for the GDS4824 dataset. (A) Accuracy. (B) AUROC.

**Fig 6.**
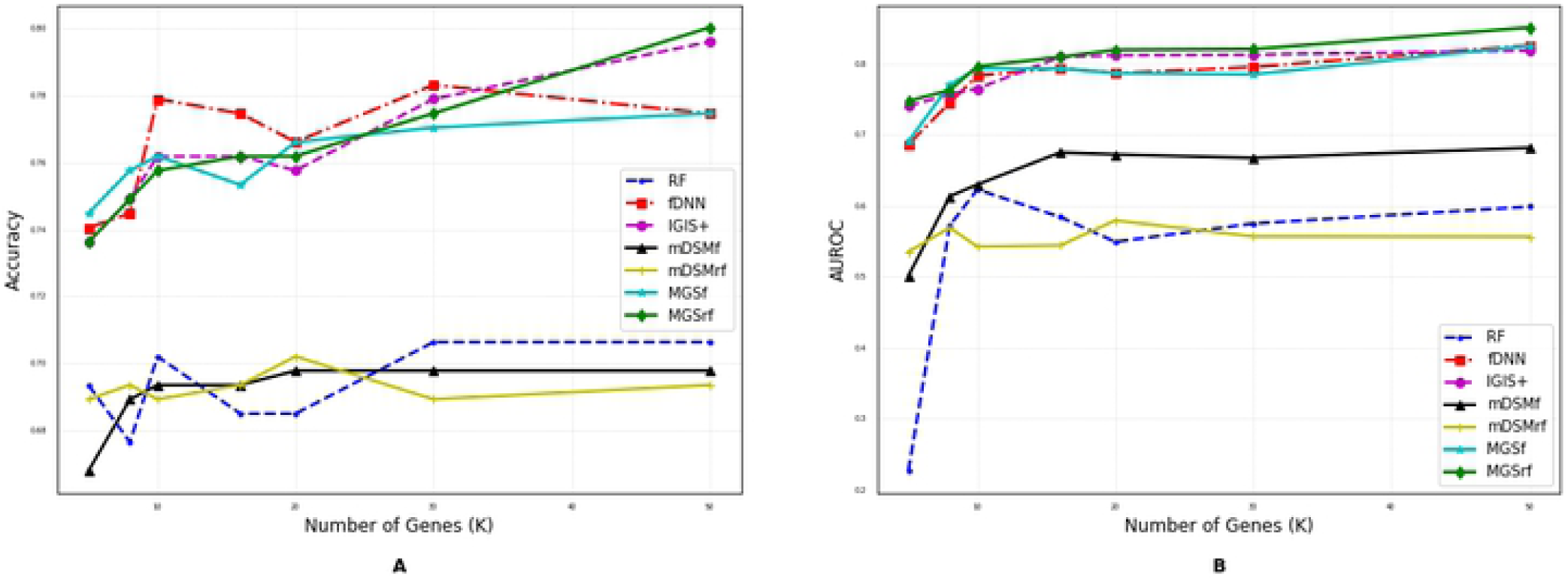
Performance comparison using different number of selected genes for the GSE106291 dataset. (A) Accuracy. (B) AUROC.

Based on the results presented in Figs 3 - 6 and Table 2, our proposed methods, *MGS*_*f*_ and *MGS*_*rf*_ outperformed the existing methods for most of the cases. The proposed filter method (*MGS*) performed well for all classifiers and thus, it is classifier independent. The datasets used for experimentation had a highly imbalanced distribution of the classes. Despite this, the performance of *MGS* was relatively better compared to the other reported methods, which also indicates that the proposed method is tolerant to the imbalanced dataset. Moreover, for every value of *η, MGS*_*rf*_ classified few more samples accurately than *MGS*_*f*_ using SVM and RF classifiers, which indicates that *MGS*_*rf*_ achieved slightly better performance.

## Conclusion

In this paper, we presented a gene selection method and two gene ranking methods for classifying high dimensional low sample size gene expression data. The proposed gene selection method utilizes the maximum relevance and complementary information for selecting informative genes that have biological importance. Experimental results on real datasets illustrate that our gene selection method consistently yields higher classification accuracy and select more biologically relevant genes than prior state-of-the-art methods do. However, there are a few challenges that are left to be addressed for further studies. First, we believe introducing higher-order gene interaction term will help to reduce the number of selected genes. Second, to obtain globally optimum gene subsets, we may need a semi-definite programming based search strategy instead of using a *χ*^2^ based filter method used in this paper.

